# White syndrome dynamics in *Porites cylindrica*: interactions among eutrophication, host structure, and microbial communities

**DOI:** 10.64898/2026.02.04.703833

**Authors:** Ewelina Rubin, Laurie Raymundo, Héloïse Rouzé

## Abstract

Coral white syndromes are among the most prevalent diseases affecting Indo-Pacific reefs, yet their etiology remains poorly understood, particularly in relation to eutrophication and microbial dynamics. Here, we investigated the drivers of *Porites cylindrica* white syndrome (PCYLWS) across two reef sites in Guam differing in nutrient enrichment and disease pressure. We combined 14 years of long-term disease monitoring with short-term lesion tracking, microbial community profiling of coral tissue and surrounding environments, and measurements of nutrient and environmental conditions. Long-term prevalence of PCYLWS was consistently higher at the oligotrophic site (Luminao) than at the eutrophic site (Tumon Bay), largely due to differences in host size structure and colony abundance. In contrast, lesions at Tumon Bay were larger and rapidly colonized by turf algae, whereas lesions at Luminao progressed faster but stabilized at smaller sizes. Microbial communities were strongly structured by sample type, with distinct assemblages in seawater, sediment, healthy tissue, and diseased tissue. Diseased tissues exhibited high spatial and temporal variability and signatures of microbial dysbiosis, characterized by reduced dominance of *Parendozoicomonas* and increased relative abundance of opportunistic taxa, including *Ruegeria*, *Muricauda*, and *Vibrio*. Environmental microbial communities and nutrient concentrations varied seasonally, particularly at the eutrophic site, suggesting transient environmental reservoirs of opportunistic bacteria. Together, these findings indicate that PCYLWS is shaped by context-dependent interactions among host population structure, environmental conditions, and dynamic microbial communities, supporting a multi-etiology framework for coral white syndromes.

**Importance:** Coral white syndromes are among the most common yet least mechanistically resolved coral diseases worldwide. By integrating 14 years of disease monitoring with lesion-scale dynamics, environmental microbiomes, and nutrient data, this study demonstrates that white syndrome expression in Porites cylindrica is shaped by context-dependent interactions among host population structure, eutrophication, and microbial dysbiosis rather than a single causative pathogen. These findings support a multi-etiology framework for coral disease and highlight the importance of environmental microbial reservoirs in modulating disease outcomes. This work provides a long-term, field-based perspective that is rarely available for coral disease systems and is broadly relevant to disease ecology in changing coastal environments.

## Introduction

Coral reefs worldwide are being lost at unprecedented rates due to multiple anthropogenic stressors, including rising seawater temperatures, overfishing, coastal development, and pollution. One of the significant threats to coral reef ecosystems is nutrient enrichment from land-based runoff and the discharge of partially treated sewage effluent (1, 2). Coral mortalities resulting from massive bleaching events and disease outbreaks are often exacerbated by coastal eutrophication, which compromises coral health and immunity, reducing their resistance to stress (3–5). On severely impacted reefs, coastal eutrophication leads to an ecological shift from coral to macroalgal-dominated ecosystems (6, 7). Higher algal abundance within the coral reef ecosystem leads to increased bacterial abundance and changes in bacterial communities via a process described as microbialization (8). During microbialization, an increase in mostly copiotrophic bacteria is supported by additional dissolved organic carbon (DOC) released by macroalgae (8, 9). Copiotrophic bacteria are more likely to be harmful to corals and cause diseases, thus giving the algae an additional competitive advantage (8, 9).

Coral disease prevalence, i.e. the percent of coral individuals affected by a disease, as well as severity, i.e. the percent of a coral colony impacted by a disease, are both increased by eutrophication in multiple regions around the world (10–13). In addition, eutrophication may lead to the emergence of new coral diseases, resulting from the increased bacterial abundance in the water column and surrounding sediment of coral reefs (4, 10, 14). Recent studies on bacterial communities in seawater surrounding coral reefs have revealed significant changes resulting from increased nutrient loads, with some bacterial taxa (e.g. *Flavobacteriaceae*) identified as indicators of ecosystem degradation (15, 16). Because the health and immunity of the coral holobiont depend on its microbiome, any major shift in the surrounding bacterial community in the water column may alter the composition of bacteria in the coral’s protective mucus layer, thereby affecting the coral’s resilience to stress and its susceptibility to disease. Therefore, eutrophication may also lead to more severe microbial dysbiosis, characterized by a shift from a healthy microbiome to a pathobiome, defined as a group of bacteria that are harmful and disease-causing (17). Monitoring changes in bacterial communities surrounding coral reefs can provide essential insights into potential sources of shifts in the coral microbiome.

Consistent with global patterns, Guam’s reefs are experiencing coral decline due to both global and local anthropogenic pressures, while also serving as a critical economic resource. Over the past two decades, scientists from Guam have reported a 37% to 60% decline in live coral cover, primarily attributed to massive consecutive bleaching events between 2013 and 2017, which were exacerbated by local environmental stressors (18, 19). In addition to bleaching mortality, Guam corals suffer from a variety of diseases, including white syndromes (WS), which are the most prevalent coral diseases on the island (20). In general, WS refers to a group of Indo-Pacific coral diseases with different etiologies but a common primary sign: tissue loss without a colored band, resulting in a white, exposed skeleton (21). WS leads to partial or whole colony mortality (22, 23). The bare white skeleton resulting from tissue loss eventually gets colonized by other encrusting organisms, primarily turf algae or cyanobacteria (24).

Guam’s coral reefs are taxonomically diverse but are dominated by *Porites* spp, including *Porites cylindrica* (25). Since host abundance is positively correlated with total disease prevalence, *Porites* spp. are disproportionately affected by multiple diseases, with five of the six observed diseases impacting this genus. *P. cylindrica* is notably the species most frequently affected by the WS lesions (20). The severity of *P. cylindrica* white syndrome (PCYLWS) is dependent on the level of dissolved inorganic nitrogen (DIN) derived from both direct and indirect human activities (12). Specifically, Tumon Bay, which is bordered by the most urbanized land on the island, exhibits low water quality and high PCYLWS severity. In contrast, areas with the lowest nitrogen enrichment, such as Luminao reef, are associated with significantly reduced disease severity (12). The difference in disease dynamics at the two reef sites is partially explained by the abundance of *Porites*, which is higher at Luminao than at Tumon Bay (20). However, the possibility of different etiologies of WS at the two contrasted N-enriched sites cannot be excluded. In addition, different nutrient conditions may influence the composition of the PCYLWS pathobiome. To date, no studies have examined the role of bacteria in the causation and progression of PCYLWS, or the effect of eutrophication on the bacterial communities associated with PCYLWS lesions.

In response to increasing anthropogenic pressure on Guam’s reefs, which includes coastal and upland development, local environmental organizations and managers are increasingly concerned about the impacts on nearshore ecosystems. The temporal and spatial variability of PCYLWS, along with the limited data on environmental factors that determine disease occurrence, pose significant challenges for forecasting future disease outbreaks. Although a growing body of evidence links nutrient enrichment to coral disease outbreaks, few studies have integrated regular *in situ* monitoring to capture the natural and seasonal variability of coral-associated microbiomes and their environmental drivers. Here, we address this knowledge gap by conducting an *in situ* field-based survey of *Porites cylindrica* White Syndrome (PCYLWS) dynamics across two reef sites with contrasting levels of eutrophication and disease pressure. By repeatedly sampling individually tagged colonies, we track lesion development and changes in the microbial community over time, enabling detection of temporal patterns often missed by snapshot surveys. In parallel, we monitor the microbial assemblages of seawater and sediment surrounding affected colonies to identify potential environmental reservoirs of opportunistic or pathogenic bacteria. Our design combines host-associated and environmental microbiome profiling with nutrient enrichment metrics, such as chlorophyll-a and nitrate concentrations, thereby providing an integrated view of the microbial ecology of coral disease under natural and anthropogenic forcing. To ground-truth our approach, we present longitudinal prevalence data from a 14-year-long-term monitoring program. This study thus provides a baseline for understanding microbial indicators of reef health and the mechanisms underlying environmentally mediated disease emergence.

## Materials and Methods

### 1. White Syndrome Dynamics

#### Long-term prevalence patterns of PCYLWS

Five reef flat coral communities along Guam’s western coast have been monitored since 2011. Three 20m x 1m permanent belt transects were established at each of the five long-term monitoring (LTM) sites: Tanguisson, Tumon Bay, West Agaña Bay, Piti Bomb Holes, and Luminao, and censused two to four times per year. At each census, all coral colonies were counted, identified to species, assigned to one of six size classes based on maximum diameter (1-10cm; 11-30cm; 31-60cm; 61-100cm; 1-2m; and >2m), and visually inspected for signs of disease and other health impacts. Diseases were identified based on (26). Prevalence per species was calculated using the following formula:

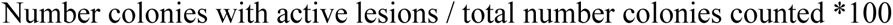

A short-term (one-year) observation of lesion dynamics and quantification of microbial community composition were conducted at two LTM sites: Tumon Bay and Luminao. The long-term PCYLWS prevalence trends are reported for these two sites. Tumon Bay is a broad, shallow embayment that was declared a Marine Preserve in 1997 and is bordered by a highly developed coastline, the focus of Guam’s tourism industry. In contrast, Luminao reef is a shallow, exposed fringe reef that developed post-WWII after the construction of the Glass Breakwater, which encloses Guam’s Apra Harbor. It is fished, but relatively inaccessible to human use.

#### Temporal and spatial variability of PCYLWS lesions

Four colonies of *P. cylindrica* were selected haphazardly at each site and tagged to monitor the dynamics of PCYLWS lesions over time. Four time points were selected, including one for the dry season (February 2024), one for the wet season (August 2024), and two during the transitional periods: May 2024 for the dry-to-wet transition and November 2023 for the wet-to-dry transition. All surveys were conducted while snorkeling. At each point, lesion monitoring and sampling were performed as follows: Day 1, five active lesions per coral colony (20 lesions per site) were identified, marked with zip ties, and photographed with a ruler for scale; Day 3, the lesions were re-photographed to document progression; and Day 5, lesions were photographed again and sampled for microbiome analyses (details below). Additionally, on Day 5, three micro-fragments were collected from visually healthy (i.e., asymptomatic) branches remote from active lesions on each tagged colony, serving as a reference for healthy tissue status. Lesions were initially scored as actively progressing if tissue loss was recent and visible as an eroding margin between dead skeleton and healthy tissue, often accompanied by mucus production within the diseased tissue zone. The sizes of the tagged lesions were measured in ImageJ^©^ by tracing their circumferences. Daily disease progression rates were calculated as:

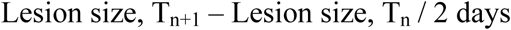

and

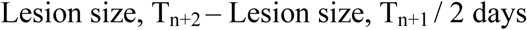

The mean progression rate was then computed from the two rates obtained for Days 3 and 5. The lesions were classified as active if the progression rate exceeded 0 cm² day^-1^, as static if the lesion area remained unchanged throughout the sampling period. None of the measured WS lesions showed a progression rate < 0, i.e., healing within the 5 days between measurements. Since the assumptions of normality and homoscedasticity were not met for the lesion size and progression rate data, we applied a non-parametric factorial analysis using the Aligned Rank Transform (ART) method, as implemented in the ARTool R package (27). This approach enabled us to test the effects of site, time point, and colony, and all interactions among these variables (site*time, site*colony, time*colony, site*time*colony) on both lesion size and progression rate without assuming normality. Pairwise comparisons were assessed using estimated marginal means with the emmeans R package (28).

### 2. Microbial communities in the environment and coral tissue

#### Sample collection

During the one-year monitoring of lesion dynamics, environmental eDNA and coral samples were collected at each of the four time points. Seawater and sediment samples were collected as follows. Three 2L seawater samples were collected and kept in coolers with ice until they reached the lab within 1 hour of collection. Seawater was filtered using sterile 0.2 µm mixed-cellulose-ester filters (Whatman; 47 mm diameter) and stored in 2 mL of RNA/DNA shield (Zymo Research) at −80 °C until DNA extraction. Sediment samples were collected in 15 mL Falcon tubes (in triplicate) and transported on ice. They were then stored frozen at −80 °C until DNA extraction. For coral samples (both remote healthy: n=3, and diseased: n=5 per colony), small fragments (∼1 cm²) of coral were cut using sterilized wire cutters. These cutters were cleaned between samples using a 10% bleach solution and 95% ethanol. Samples were preserved in an RNA/DNA shield (Zymo Research), stored in a cooler with ice during field work and transport, and frozen at −80 °C upon arrival at the Marine Laboratory.

#### DNA extractions and V4-rRNA libraries

The microbial (bacterial and archaeal) community compositions of healthy and diseased coral tissue, as well as those of eDNA (seawater and sediment) samples, were determined using the variable region 4 (V4) of the 16S rRNA gene amplicon via Next-Generation Illumina sequencing. First, DNA was extracted using the Zymo BIOMICS DNA Extraction Kit (Zymo Research). PCRs for the V4 region of the 16S rRNA gene were amplified using modified forward 515F (29) and reverse 806R (30) primers. The purified product from the first PCR (GeneJet; Thermo Fisher©) was used in the second PCR, which was designed to add Illumina adapters and the 6-mer dual indices, as previously described (31). Both PCRs were generated using Takara PCR buffer and Hot Start polymerase (Takara). The first PCR was carried out in 30 µl reactions with the following final concentrations: dNTPs at 0.8 mM, each primer at 0.5 µM, and 30 to 100 ng of total DNA. The first PCR cycling conditions were as follows: a hot-start activation step for 2 min at 95 °C, followed by 30 cycles of 95 °C for 40 s, 58 °C for 2 min, and 72 °C for 1 min, with a final extension of 5 min at 72 °C. The second PCR was performed in a 20-µl reaction volume using the same components and final concentrations as above, with 30 ng of the PCR-1 product serving as the template. The second PCR cycling condition was as follows: a hot-start activation at 95 °C for 2 min, followed by five cycles of 95 °C for 40 s, 59 °C for 2 min, and 72 °C for 1 min, and a final extension at 72 °C for 7 min. All samples were pooled and gel-purified using the Qiagen Gel Purification Kit. The V4-16S library pool was sent to CD Genomics (US), a commercial sequencing company for Illumina MiSeq sequencing, using a 2 x 300 bp cycling kit. After downloading the data, the quality of the raw sequencing reads for each sample was evaluated using FASTQC.

#### Microbiome/Bacterial data analysis

Data analyses were conducted in R v. 4.5.1 using multiple packages, including *DADA2* (32), *phyloseq* (33), *microbiome* (34), and *microViz* (35). After quality inspection, the raw reads were trimmed and filtered, using a 260 bp cutoff for both forward and reverse reads. A minimum quality score of 10 or higher was set using the DADA2 package. The same package was used to generate the Amplicon Sequences Variants (ASV) sequence table and to remove chimeric ASVs. Taxonomy was assigned using the SILVA SSU rRNA database v.138.2 (36). ASVs classified as mitochondria, chloroplasts, eukaryotes, or an unknown Kingdom were removed from the final count table. Additionally, only ASVs with an average of five reads per sample were retained for the final count table. Count data were transformed to relative abundance and visualized using stacked bar charts constructed in the library *fantaxtic* (37). All other plots were constructed using the *ggplot2* (38). The Bray-Curtis dissimilarity between samples was calculated, and the resulting dissimilarities were visualized using Principal Coordinates Analysis (PCoA) with the *microViz* package. A permutational multivariate analysis of variance (PERMANOVA) was conducted to test the difference in beta diversity using the same package. Pairwise Adonis was performed using the package *pairwiseAdonis* (39). The microbial bacterial communities were first compared among the four sample types: seawater, sediment, healthy coral tissue, and diseased coral tissue. Because the samples from different origins (sediment, seawater, healthy, and diseased tissue) had distinct microbial communities, the dataset was separated by sample type for further analyses to examine differences between sites and time points. Differential abundance (DA) analysis was performed using the beta-binomial regression model and either the Wald test for a single variable or the Likelihood Ratio test for multiple variables, both implemented in the *corncob* package (40). Pearson’s correlation coefficients were calculated to assess relationships between genus-level relative abundance and both lesion size and progression rates, to identify the highest contributors to coral tissue loss.

### 3. Environmental Monitoring

Chlorophyll *a* concentration of seawater (as a proxy of phytoplankton biomass, and thus also nutrient load) was measured through three 1-L seawater samples (in triplicate) collected at the two reef sites every other month for one year, from October 2023 to August 2024. For the nitrate and ammonium concentrations, 50 mL of seawater samples (in triplicate) were collected at the two reef sites every other month for the same period. All samples were transported to the lab in coolers with ice for further processing. Seawater samples were then filtered using GF-F filters (Whatman, 0.7 μm pore size, 0.47 mm diameter). Filters were then placed into 5 mL of 100% acetone and incubated for 24 hours at 4 °C, protected from light exposure. Chlorophyll *a* concentration was measured using the Trilogy Laboratory spectrophotometer Fluorometer (Turner Designs) with the Chl a non-acidified module. The seawater samples for nitrate analysis were stored at −80 °C until analysis. Nitrate analysis of seawater samples was conducted using the LaMotte Test Kit Method (Trilogy, Turner Designs) and the nitrate 540 nm absorbance module on the Trilogy Laboratory Fluorometer (Turner Designs) using the Nitrate absorbance module. An ammonium analysis was carried out using the method described by (41). All ammonium readings were below the detection limit of 0.1 mg/L; thus, the data are not presented. Temperature loggers (Onset^©^ HOBO tidbits) were deployed at both reef sites as part of the long-term monitoring program for coral health and disease, as described above. Precipitation data for Guam were obtained from the NOAA National Weather Service website (https://www.weather.gov/)

## Results

### 1. White Syndrome Dynamics

#### Long-term prevalence patterns of PCYLWS

PCYLWS prevalence differed markedly between Luminao and Tumon Bay over the 2011–2025 monitoring period, with Luminao consistently exhibiting higher disease levels than Tumon Bay across all colony size categories (Figure 1A). Size-stratified analyses revealed a clear size-dependent pattern at both sites, with WS prevalence increasing with colony size (Figure 1A). At Luminao, transect-weighted mean annual WS prevalence calculated across years increased from 6.36% in colonies <10 cm in diameter to 59.13% in colonies >2 m (Figure 1A). Tumon Bay exhibited substantially lower WS prevalence across all size classes, with most disease occurrences concentrated in intermediate and large colonies (61–100 cm) and transect-weighted annual WS prevalence across years reaching 27.28% in colonies >1 m in diameter (Figure 1A). Notably, colonies >2 m in diameter—those exhibiting the highest WS prevalence at Luminao—were absent from Tumon Bay (Figure 1A, B), likely contributing to the lower overall disease prevalence observed at this site.

**Figure 1.**
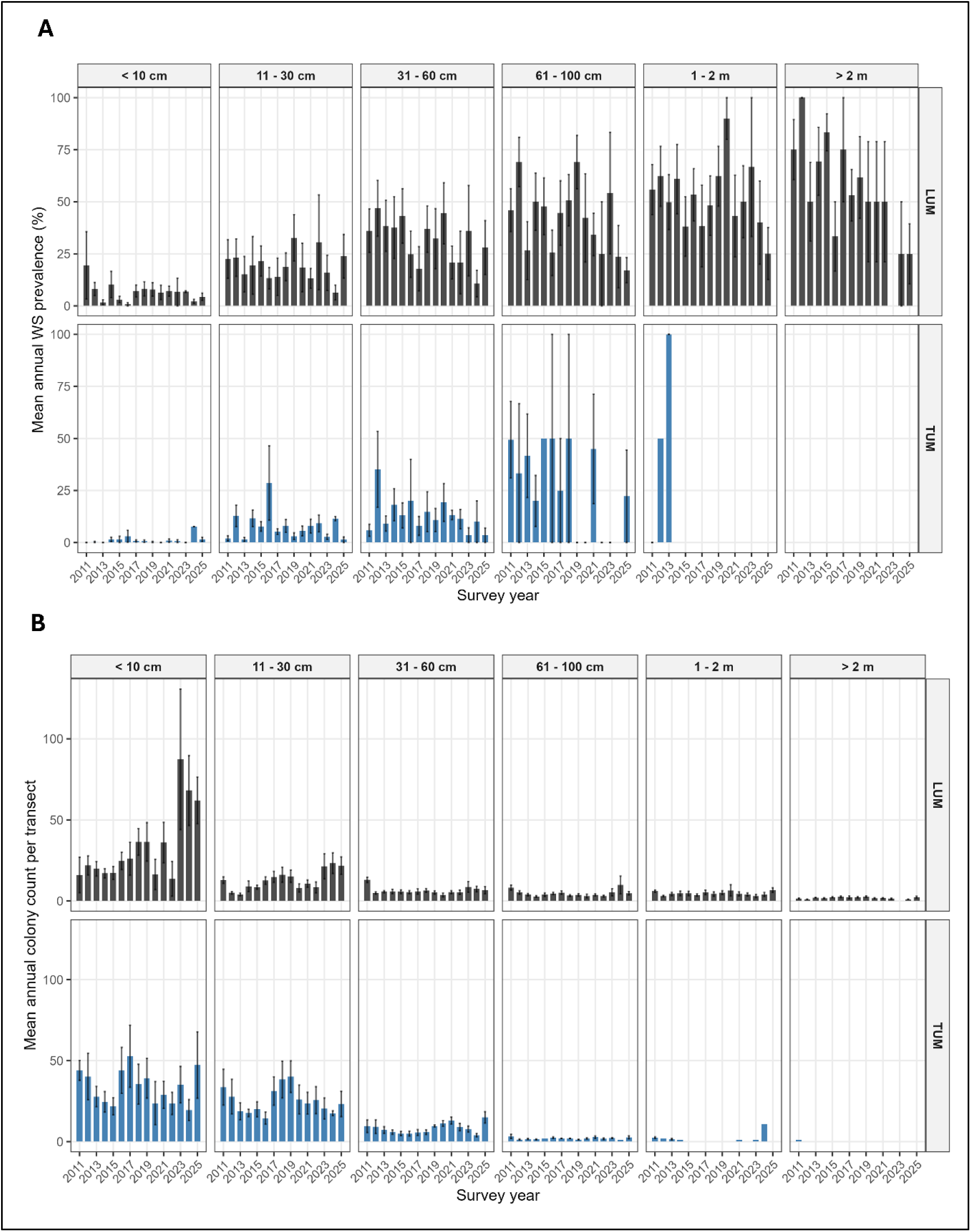
Long-term (14 years) record of *Porites cylindrica* white syndrome (PCYLWS) at two reef sites. Annual size-dependent white syndrome prevalence (A) and coral colony abundance (B) at Luminao (grey) and Tumon Bay (blue) between 2011 and 2025. WS prevalence was calculated per transect and averaged annually; colony abundance represents mean transect-level counts. Error bars denote ±1 standard error. Size classes are ordered from smallest to largest across the panel.

#### Temporal and spatial variability of PCYLWS lesions

There was considerable variation in the gross appearance of PCYLWS lesions across the two study sites during lesion monitoring between November 2023 and August 2024 (Figure 2C-D). Lesion sizes differed significantly between sites (ART test; F = 42.07, p < 0.001) and among time points (ART test; F = 4.14, p = 0.007), but there was no significant interaction between site and time point (ART test; F = 2.40, p = 0.07). Coral lesion size in Tumon varied from 2.18 ± 1.56 cm^2^ (mean ± SD) in February 2024 to 4.81 ± 5.04 cm^2^ in August 2024 (Figure 2A). On the other hand, lesions in Luminao were smaller, ranging from 0.69 ± 0.43 cm² in May 2024 to 1.30 ± 1.14 cm² in August 2024 (Figure 2A). There was no difference in lesion size among different colonies in either Tumon (ART test; F=2.49, p=0.069) or in Luminao (ART test, F=2.16, p=0.10).

**Figure 2.**
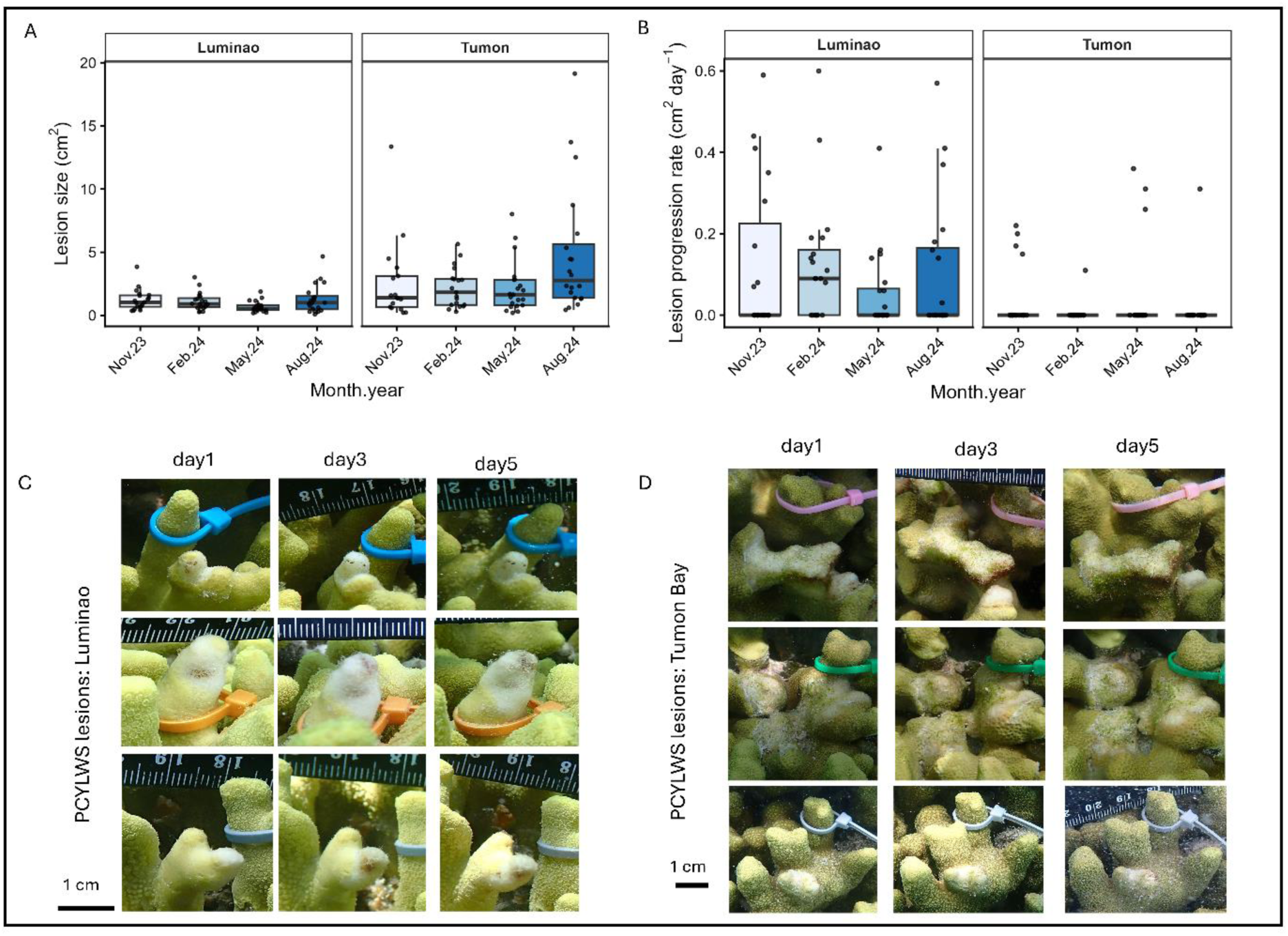
Lesion size, progression, and morphology of *Porites cylindrica* white syndrome (PCYLWS) at two reef sites. (A) Lesion size (cm²) at Luminao and Tumon Bay across four seasonal sampling periods. Boxes show medians and interquartile ranges; points represent individual lesions. (B) Lesion progression rates (cm² day⁻¹) across the same periods; zero values indicate static lesions. (C) Representative PCYLWS lesions at Luminao were photographed on Days 1, 3, and 5, typically located at branch tips and remaining small. (D) Representative PCYLWS lesions at Tumon Bay were photographed on Day 1, Day 3, and Day 5, generally larger and occurring between branches, with the exposed skeleton rapidly overgrown by turf algae. Scale bars = 1 cm.

Colonies at Luminao exhibited a higher proportion of active lesions (progression rate > 0), with 46% of lesions showing measurable progression, compared to 18% at Tumon. Lesion progression rates differed significantly between sites (ART ANOVA, F₁,₁₄₈ = 8.65, p = 0.0038), with higher progression at Luminao than Tumon. In contrast, no significant effect of month (F₃,₁₄₈ = 0.77, p = 0.51) and no site × month interaction (F₃,₁₄₈ = 0.33, p = 0.80) were detected, indicating that temporal patterns in lesion progression were consistent across sites (Figure 2B). Considering only active lesions, mean progression rates were 0.0706 cm² day⁻¹ at Luminao and 0.0225 cm² day⁻¹ at Tumon. Lesion progression rates did not differ among colonies within either Tumon (ART ANOVA, F₇,₇₀ = 0.65, p = 0.59) or Luminao (ART ANOVA, F₇,₇₈ = 0.19, p = 0.90). Healing of the lesions (via re-sheeting of new tissue and skeleton) was observed over a period of 2 to 3 months. Due to the longer timescale, the healing process was not quantified.

There were also etiological differences in lesion location on coral colonies. The PCYLWS lesions at Luminao were usually located on the tip of the branches, and tissue loss progressed from the tip down, but did not reach the base of the branch. The lesions remained small, and a rigid margin around them was frequently observed (Figure 2C). In Tumon Bay, PCYLWS lesions were larger and typically found between coral branches rather than at their tips. The dead skeleton was quickly covered by green and red turf algae (Figure 2D), with a distinct, actively progressing lesion margin. Thus, the bare skeleton did not tend to re-sheet, due to substantial algal cover (Figure 2D).

### 2. Microbial community composition in the environment and coral tissue

#### Alpha and beta diversity in the environment and coral tissue samples

A total of 604 ASVs were identified in the entire dataset, representing 27 phyla, 40 classes, 84 orders, 143 families, and 267 genera. The ASV richness of the microbial (archaeal and bacterial) community was the highest in sediment, followed by seawater and diseased tissue, and lowest in the healthy tissue (Figure 3). Alpha diversity of microbial communities in seawater and coral tissue samples (both healthy and diseased) was significantly different between the two sites (Figure 3; t-test, p < 0.05). However, there was no difference in alpha diversity in the sediment samples between the two sites (Figure 3, t-test, p > 0.05). Alpha diversity in seawater and healthy tissue was higher in Luminao samples than in Tumon samples; however, in diseased tissue, the pattern was reversed, with higher diversity in the Tumon samples (Figure 3). Beta diversity varied significantly among the four sample types: seawater, sediment, healthy tissue, and diseased tissue (Figure 4; PERMANOVA, F = 57.72, p < 0.01). All pairwise comparisons were significantly different (p_adj_ < 0.001, Supplementary File).

**Figure 3.**
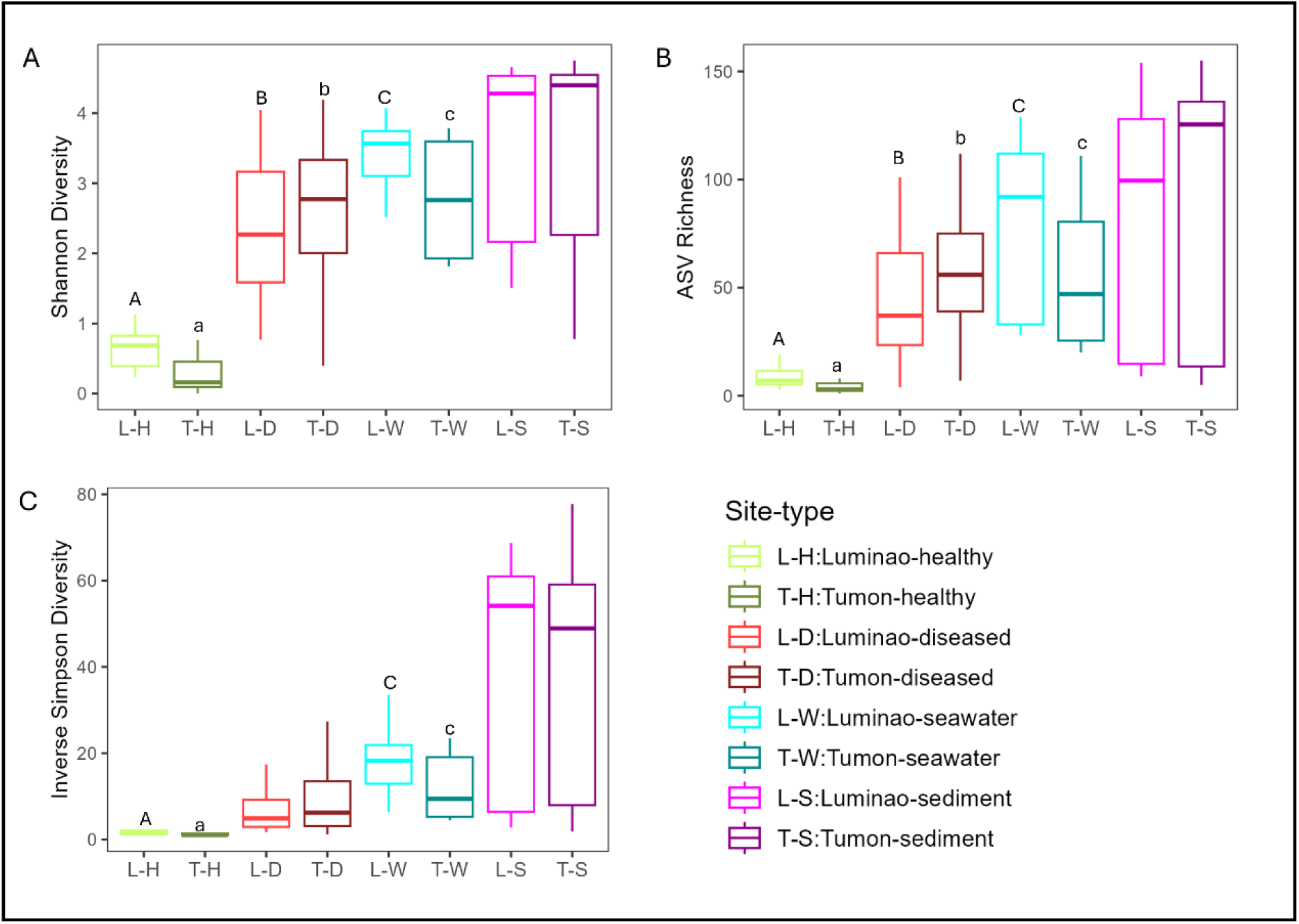
Alpha diversity of microbial communities across sites and sample types. (A)Shannon diversity, (B) ASV richness, and (C) inverse Simpson diversity of microbial communities in healthy coral tissue, diseased coral tissue, seawater, and sediment at Luminao and Tumon Bay. Boxes show medians and interquartile ranges; whiskers indicate data range. Different letters denote significant differences among sample types (p < 0.05).

**Figure 4.**
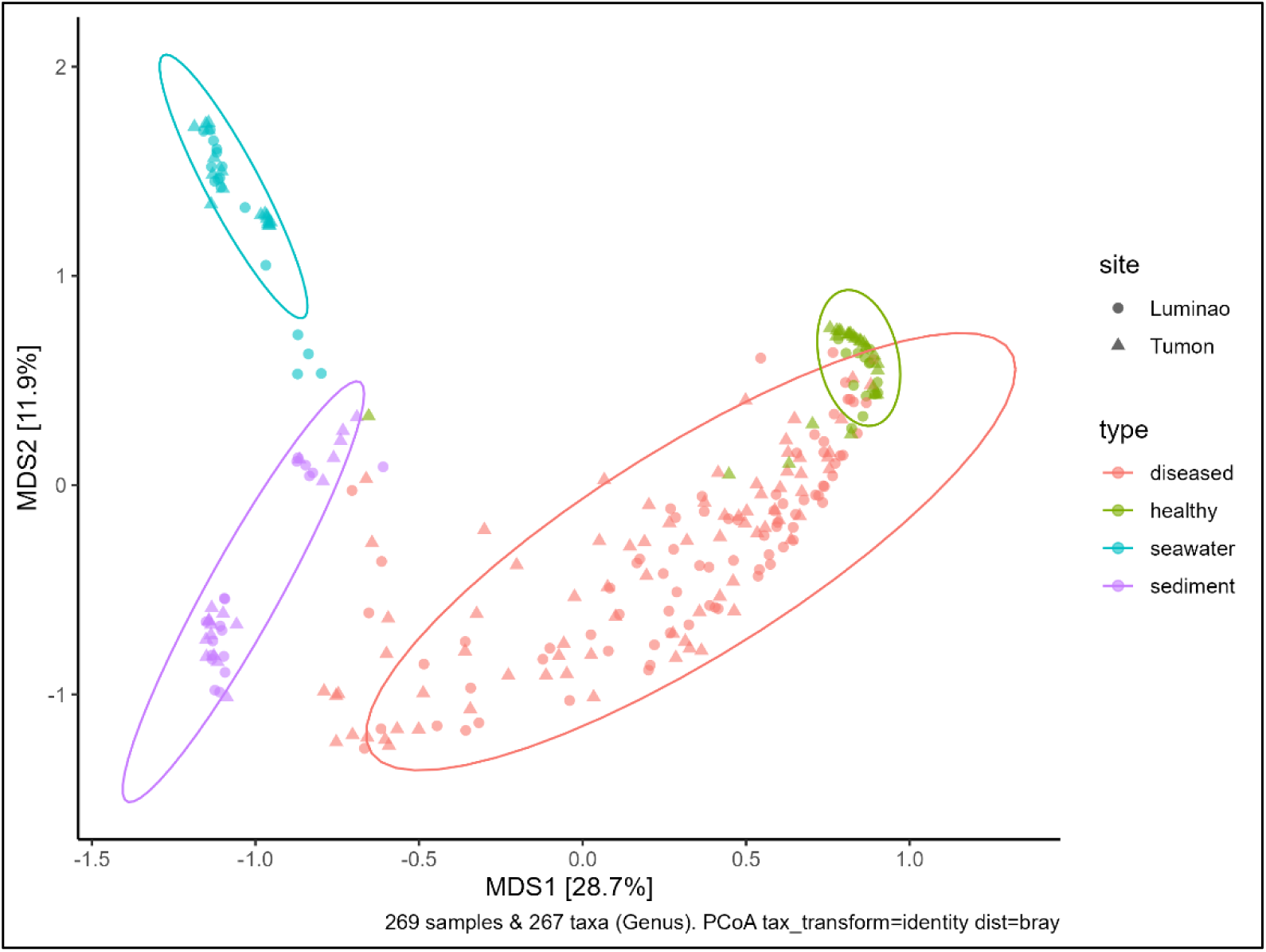
Beta diversity of microbial communities across sample types and sites. Non-metric multidimensional (NMDS) based on Bray–Curtis dissimilarity of microbial communities from coral tissue, diseased coral tissue, seawater, and sediment collected at Luminao and Tumon Bay. Points represent individual samples (n=269); colors indicate sample type and symbols indicate site. Ellipses represent group dispersion. Community composition differed significantly among sample types (PERMANOVA, p < 0.01).

#### Differential abundance among environmental and coral tissue samples

Bacterial relative abundances differed significantly among the four sample types (Figure 5). Specifically, 17 bacterial orders contributed most to these differences (Figure 5; Wald test, FDR < 0.05). *Pseudomonadales* showed the highest relative abundance in healthy tissue (92–100%) compared to diseased tissue, sediment, or seawater samples (Figure 5, Wald test, FDR < 0.05). *Flavobacteriales* (23–49%), *Enterobacterales* (6–19%), and *Verrucomicrobiales* (1–7%) exhibited higher relative abundance in seawater than in any other sample type (Figure 5, Wald test, FDR < 0.05). In sediment samples, *Desulfobacterales* (19–57%), *Cyanobacteriales* (12–35%), and *Kiloniellales* (3–8%) were among the top orders with higher relative abundances compared to seawater, healthy tissue, or diseased tissue (Figure 5, Wald test, FDR < 0.05). *Phormidesmiales* (3–41%), *Cytophagales* (4–18%), *Campylobacterales* (1–14%), and *Chitinophagales* (2–9%) were enriched in diseased tissue compared to either healthy tissue or environmental samples (Figure 5, Wald test, FDR < 0.05).

**Figure 5.**
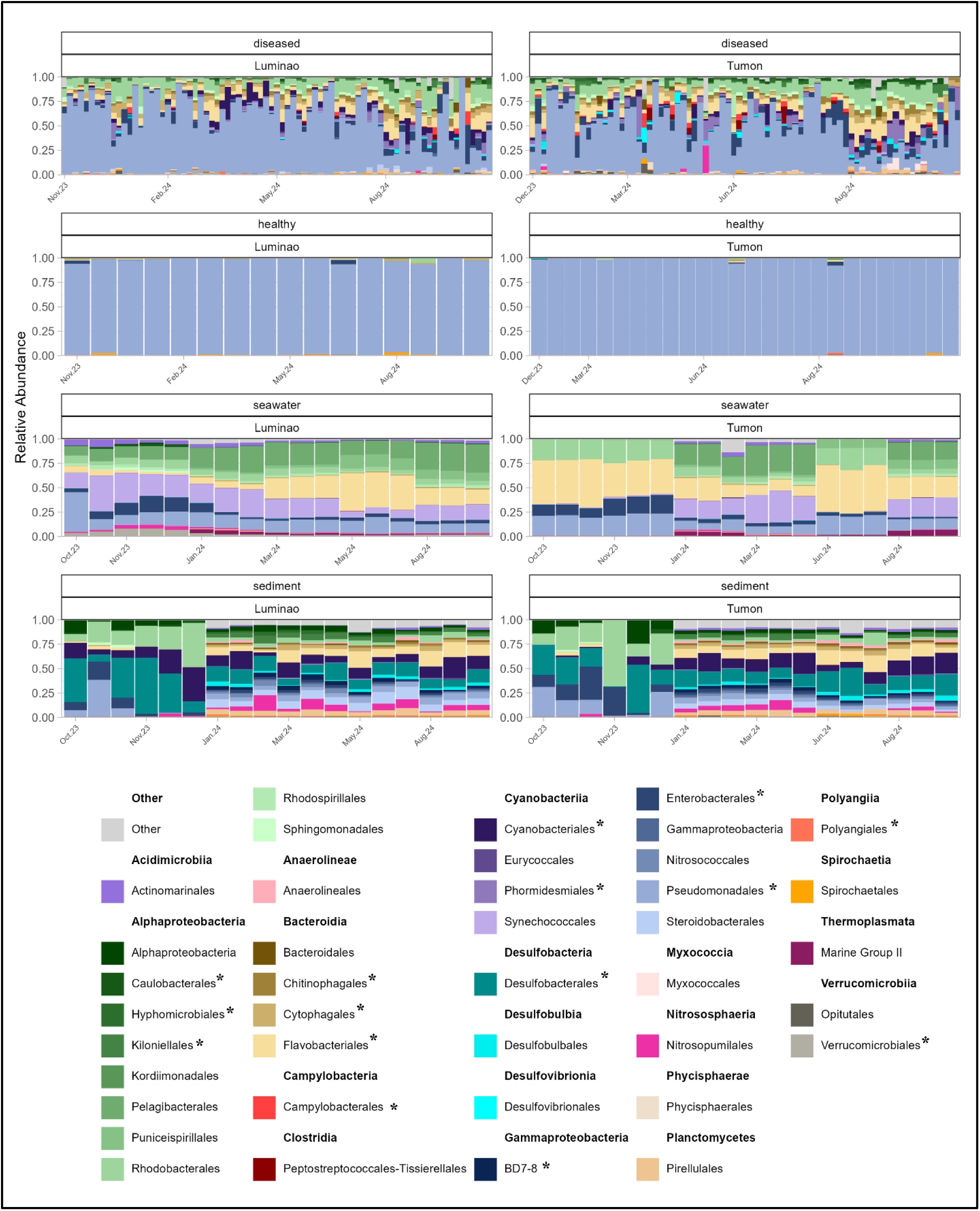
Relative abundance of the top 40 most abundant microbial orders in four sample types 1) healthy (n =40) and diseased 2) (n = 156) tissue of *P. cylindrica*, 3) seawater (n = 35), and 4) sediment (n = 36), collected from two reef sites (Luminao and Tumon) between October 2023 and August 2024. The asterisk next to the class name indicates a significant differential relative abundance between sample types (Wald test, false discovery rate [FDR] < 0.05).

#### Temporal and spatial variability of microbial communities in eDNA samples

Beta diversity of sediment samples (n = 36) from the two sites showed highly similar patterns (Figure 6A; PERMANOVA, F = 1.44, p = 0.2) but varied significantly across time points (Figure 6A; PERMANOVA, F = 9.09, p < 0.001). In particular, samples taken in October and November 2023 formed a distinct cluster from those collected in other months (Figure 6A). Among the 55 bacterial orders detected in sediment, the relative abundance of 17 orders varied by both site and time point (Supplementary File; Likelihood Ratio Test (LRT), FDR < 0.05). An additional three orders differed between sites but not across time points, while 21 orders varied over time but not between sites (Supplementary File; LRT, FDR < 0.05). Temporal patterns of the 10 differentially abundant (DA) orders with the highest mean relative abundance are shown in Figure 7. *Enterobacterales* were more abundant in sediment samples from Tumon (Figure 7D; LRT, FDR < 0.05), while *Steroidobacterales* were more abundant in Luminao (Figure 7E; LRT, FDR < 0.05). *Rhodobacterales* and *Enterobacterales* showed their highest sediment relative abundances in October and November 2023 (Figure 7A, D; LRT, FDR < 0.05), whereas *Flavobacteriales* peaked in May and August 2024 (Figure 7C; LRT, FDR < 0.05).

**Figure 6.**
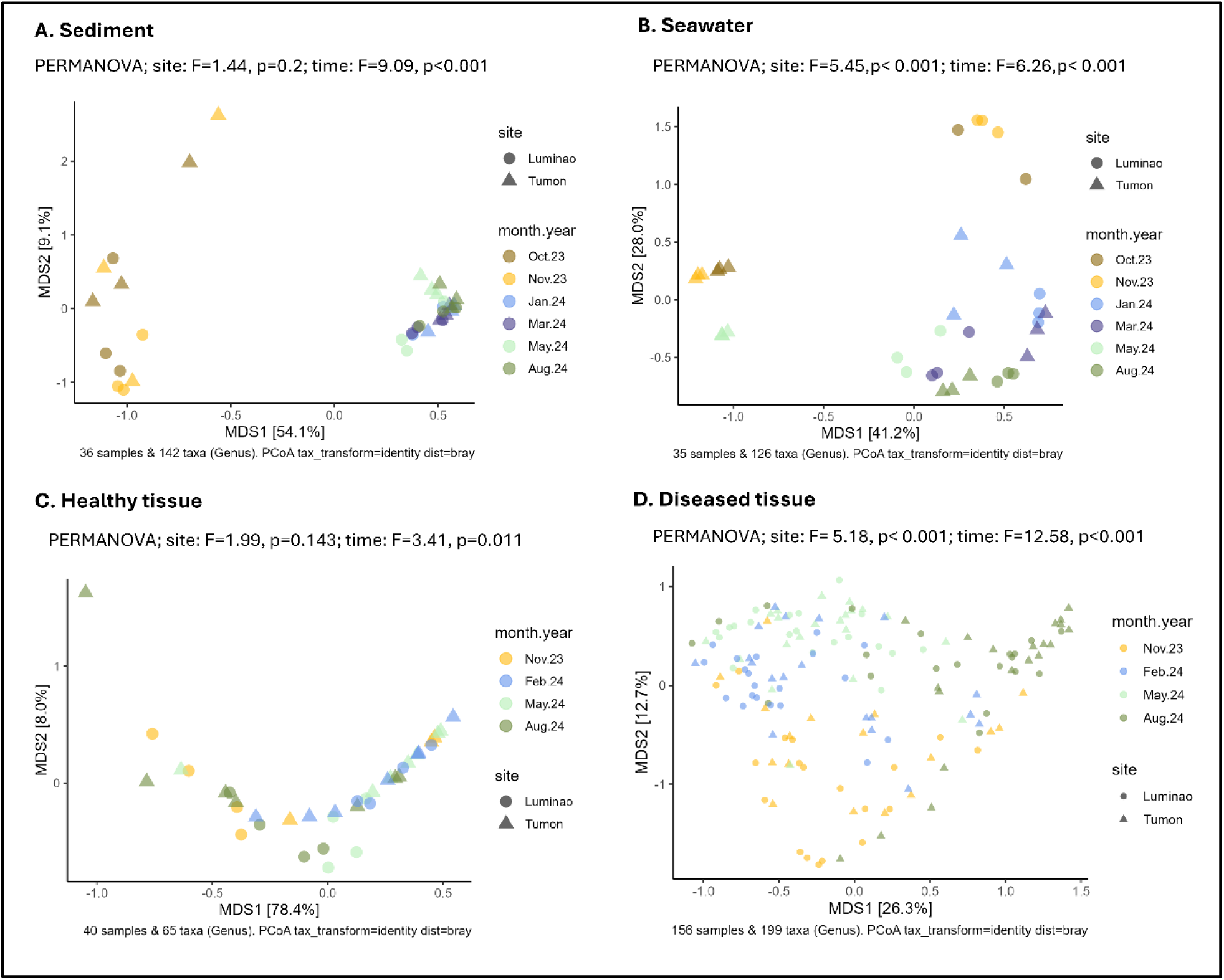
Non-metric multidimensional ordination using the Bray-Curtis dissimilarity matrix for microbial community composition generated from two reef sites and four sample types, including seawater (A), sediment (B), diseased (C), and healthy (D) coral tissue.

**Figure 7.**
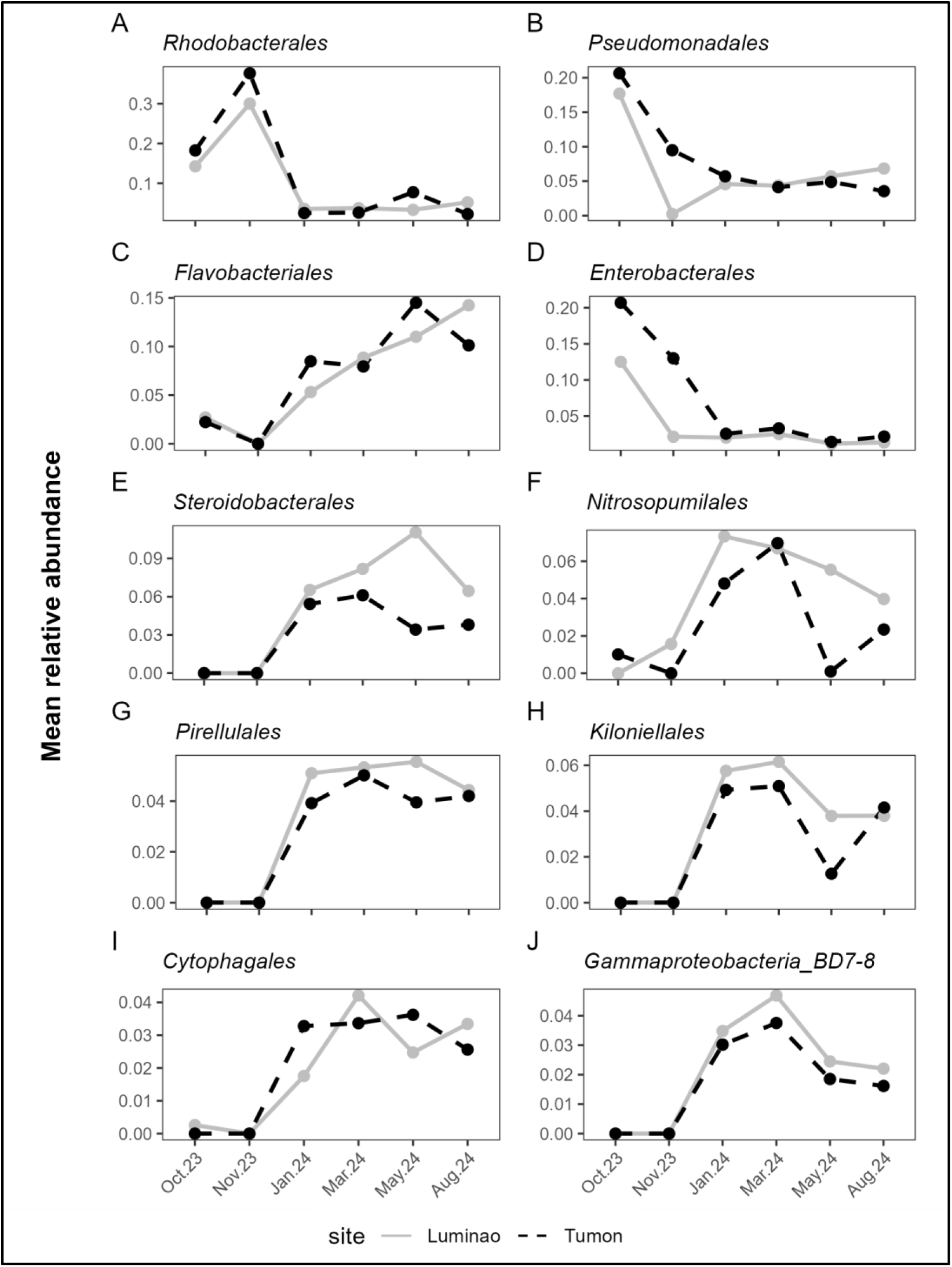
The top 10 most abundant orders in sediment samples that were significantly differentially abundant (DA) between reef sites, across time points, or both (Likelihood Ratio Test, FDR < 0.05).

Microbial communities in seawater samples (n = 35) varied significantly across both sites and time points (Figure 6B; PERMANOVA, site: F = 5.45, p < 0.001; time: F = 6.26, p < 0.001). Among the 52 microbial orders detected in seawater, the relative abundances of 35 orders varied at both spatial and temporal scales. An additional two orders differed between sites but not over time, while 16 orders varied across time points but not between sites (Supplementary File; LRT, FDR < 0.05). Among the ten most abundant orders, *Flavobacteriales*, *Rhodobacterales*, *Pelagibacterales*, and *Synechococcales* showed different temporal patterns between the two reef sites (Figure 8A, B, C, E; LRT, FDR < 0.05). In Tumon, *Flavobacteriales* and *Rhodobacterales* were enriched in October and November, and again in May (Figure 8A, E; LRT, FDR < 0.05). In contrast, their relative abundances in Luminao increased only in May (Figure 8A, E; LRT, FDR < 0.05). *Pelagibacterales* and *Synechococcales* showed relatively stable levels in Luminao throughout the study (Figure 8B, C). However, in Tumon, their relative abundances increased primarily in January, March, and again in August (Figures 8B and 8C; LRT, FDR < 0.05). *Enterobacterales* displayed similar temporal patterns at both reef sites, with higher relative abundances in October and November compared to all other months (Figure 8F; LRT, FDR < 0.05).

**Figure 8.**
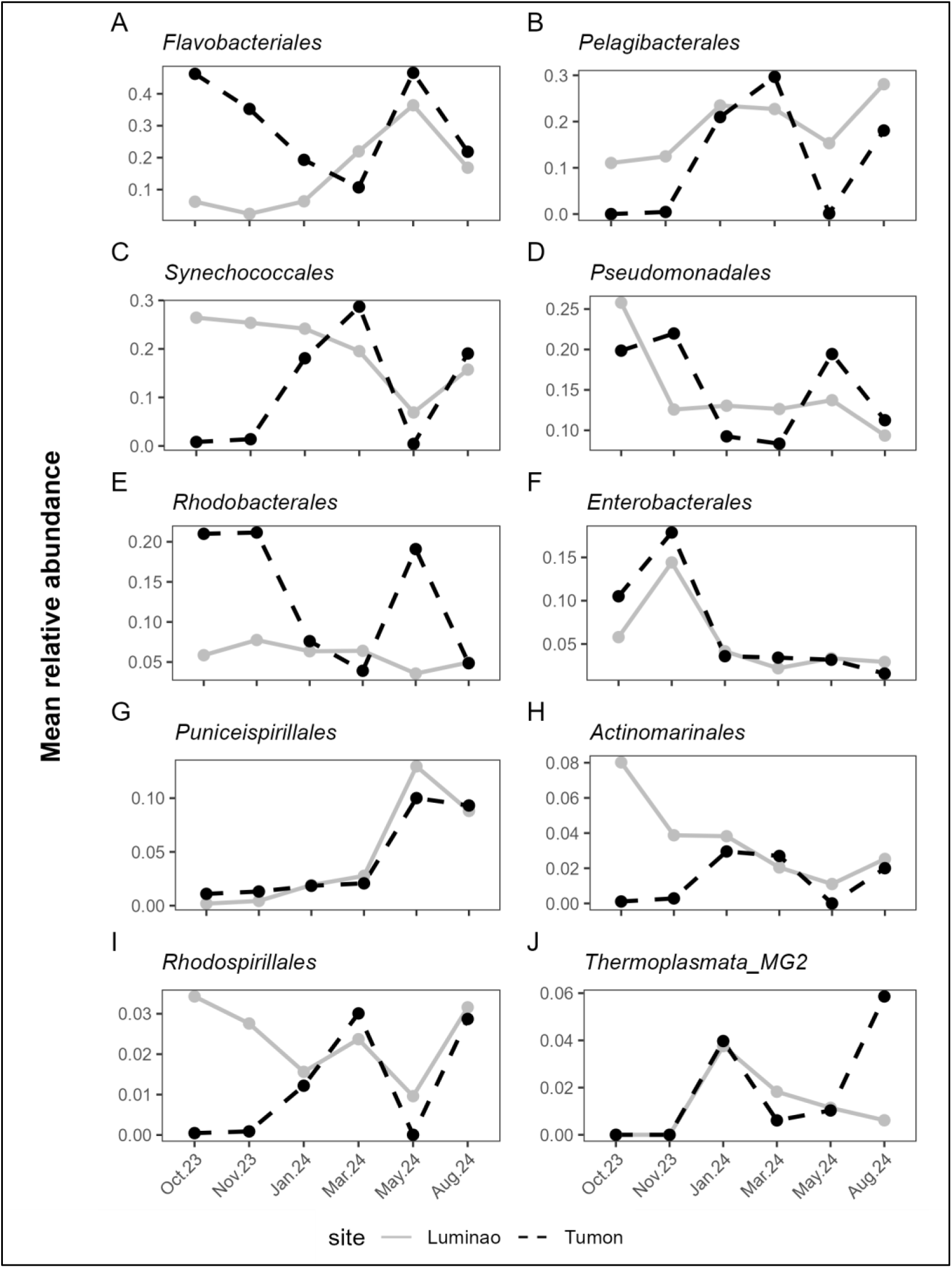
The top 10 most abundant orders in seawater samples that were significantly differentially abundant between reef sites, across time points, or both (Likelihood Ratio Test, FDR < 0.05).

#### Temporal and spatial variability of microbial communities in coral tissue samples

Healthy coral tissue samples (n = 40) showed similar microbial communities, with no significant differences in beta diversity between sites and no clear seasonal patterns across time points (Figure 6C; PERMANOVA, time: F = 3.41, p = 0.011). In contrast, the microbial community in diseased tissue samples (n = 156) differed significantly between sites and across time points (Figure 6D; PERMANOVA, site: F = 5.18, p < 0.001; time: F = 12.58, p < 0.001). Additionally, pairwise comparisons of beta diversity showed that microbial communities in diseased tissue varied significantly across all time points at both sites (Supplementary File: pairwise PERMANOVA results).

Of the 63 bacterial orders detected in diseased tissue samples, the relative abundances of 18 orders varied across both site and time point. An additional eight orders differed between sites but not over time, while 10 varied across time points but not between sites (Supplementary File; LRT, FDR < 0.05). Among the ten most abundant orders in lesions, *Rhodobacterales* contributed, on average, between 5.5% and 14.1% of the microbial community, reaching as high as 32.7% in some samples. Although its relative abundance did not differ between reef sites, it varied significantly over time, with the lowest relative abundance in February 2024 at both locations (Figure 9B; LRT, FDR < 0.05). *Phormidesmiales* and *Caulobacterales* were more abundant in lesions from Tumon at all time points (Figure 9E and 9I; LRT, FDR < 0.05). Similarly, *Enterobacterales* and *Campylobacterales* had higher relative abundances in Tumon Bay lesions at three of the four time points (Figure 9D, 9J; LRT, FDR < 0.05). In contrast, *Cyanobacteriales* were more abundant in lesions from Luminao (Figure 9F; LRT, FDR < 0.05).

**Figure 9.**
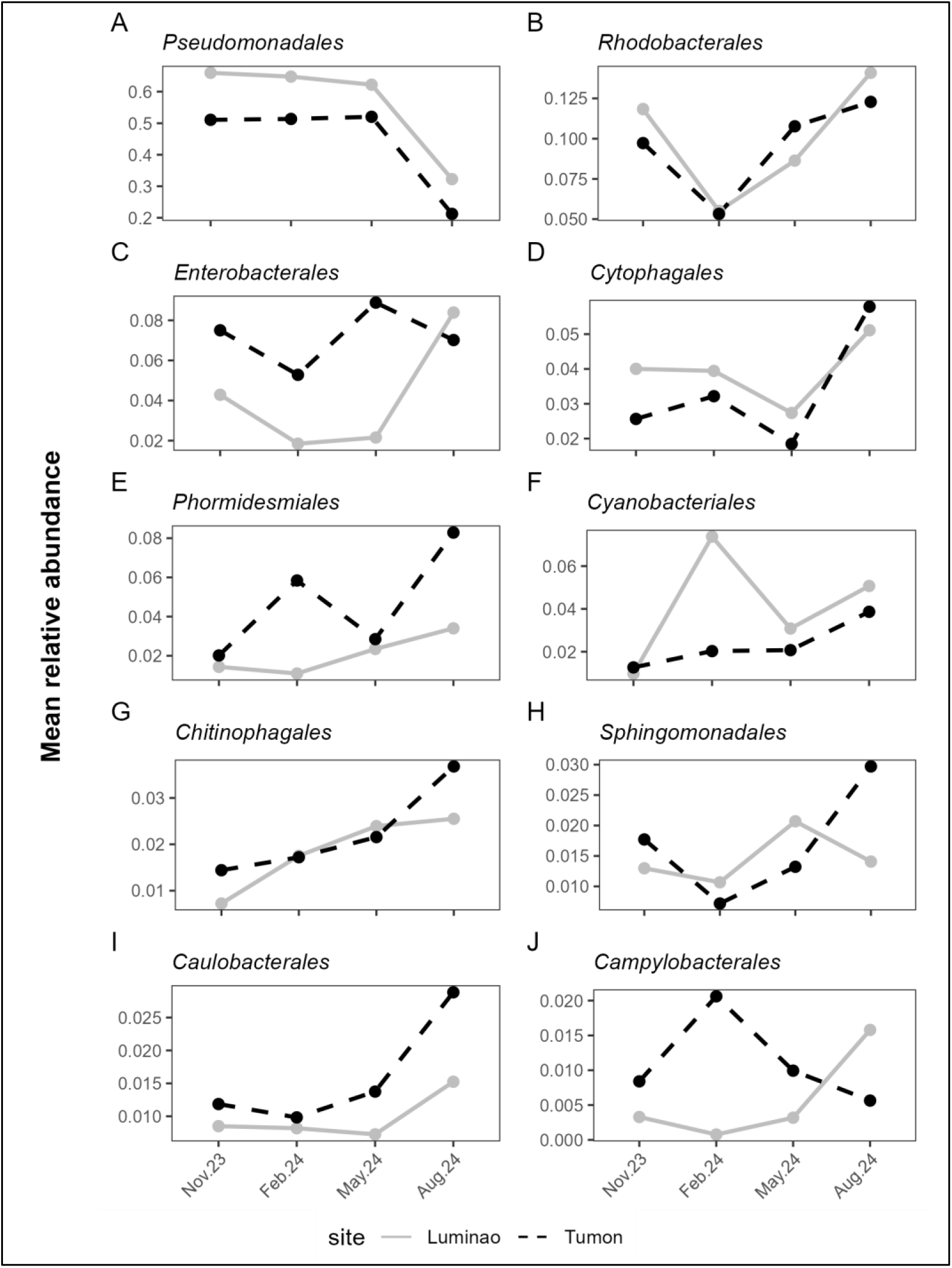
The top 10 most abundant orders in WS-affect tissue *of P. cylindrica* that were significantly differentially abundant between reef sites, across time points, or both (Likelihood Ratio Test, FDR < 0.05).

#### Differential abundance: Healthy vs Diseased Tissue

Striking differences in bacterial relative abundance were observed between healthy and diseased coral tissue. In Tumon, 190 microbial genera were detected in both healthy and diseased tissue samples. Of these, 121 were differentially abundant (DA) between diseased and healthy tissue, after controlling for sampling time and colony ID (Supplementary File; LRT, FDR < 0.05). In Luminao, 161 genera were detected, of which 80 showed differential abundance between healthy and diseased tissue (Supplementary File; LRT, FDR < 0.05). Seventy DA genera were shared between the two sites, while 51 were unique to Tumon and 11 were unique to Luminao. The 43 most abundant genera (with a minimum mean relative abundance of 0.5%) that were differentially abundant between healthy and diseased samples at either site are shown in Figure 10. The two genera with the highest prevalence in disease samples were *Ruegeria* and *Muricauda*, found in 90% and 84% of lesions in Luminao, and in 91% and 87% of lesions in Tumon, respectively (Figure 10). Additionally, *Vibrio* was detected in 43% of lesions in Luminao and 74% in Tumon (Figure 10). *Ruegeria* and *Vibrio* were also present in a substantial proportion of healthy samples: 25% and 19% in Lumimao and 15% and 12% in Tumon, respectively (Figure 10). *Muricauda* was less prevalent in healthy samples: 6% in Luminao and 4% in Tumon (Figure 10). Microbial diversity in lesions was high: between 9 and 16 genera accounted for 5% to 1% of the total microbial abundance, while over 100 genera each contributed less than 1%. *Parendozoicomonas* was the dominant genus in all healthy samples, regardless of site (Figure 10). Although its relative abundance decreased in diseased tissue, it still made up approximately half of the bacterial community (Figure 10). *Sulfurimonas* and *Fusibacter* had the highest positive correlation with lesion progression rates (Pearson’s r = 0.5 and 0.4, respectively; Figure 11). In addition, *Paucidesulfovibrio* and an unclassified genus within the family *Flavobacteriaceae* showed positive correlations with lesion size (Pearson’s r = 0.6–0.5; Figure 11).

**Figure 10.**
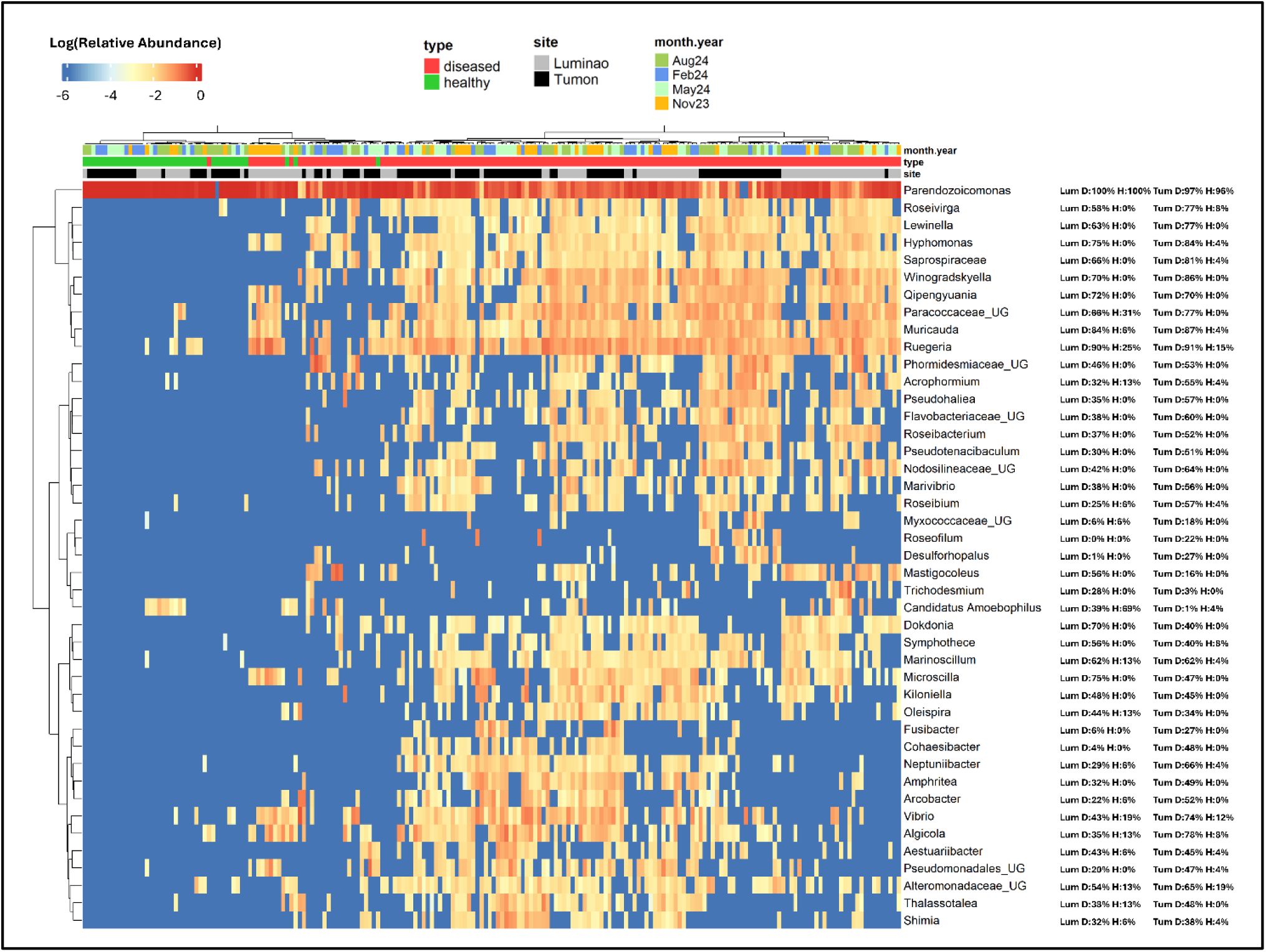
Heat map of log-transformed relative abundance (RA) for 43 genera found to be differentially abundant between healthy and diseased tissue in Luminao and Tumon samples (LRT, FDR < 0.05). The numbers in the table on the right correspond to the prevalence of that genus in diseased (D) and healthy (H) tissue samples in Luminao (Lum) and Tumon (Tum)

**Figure 11.**
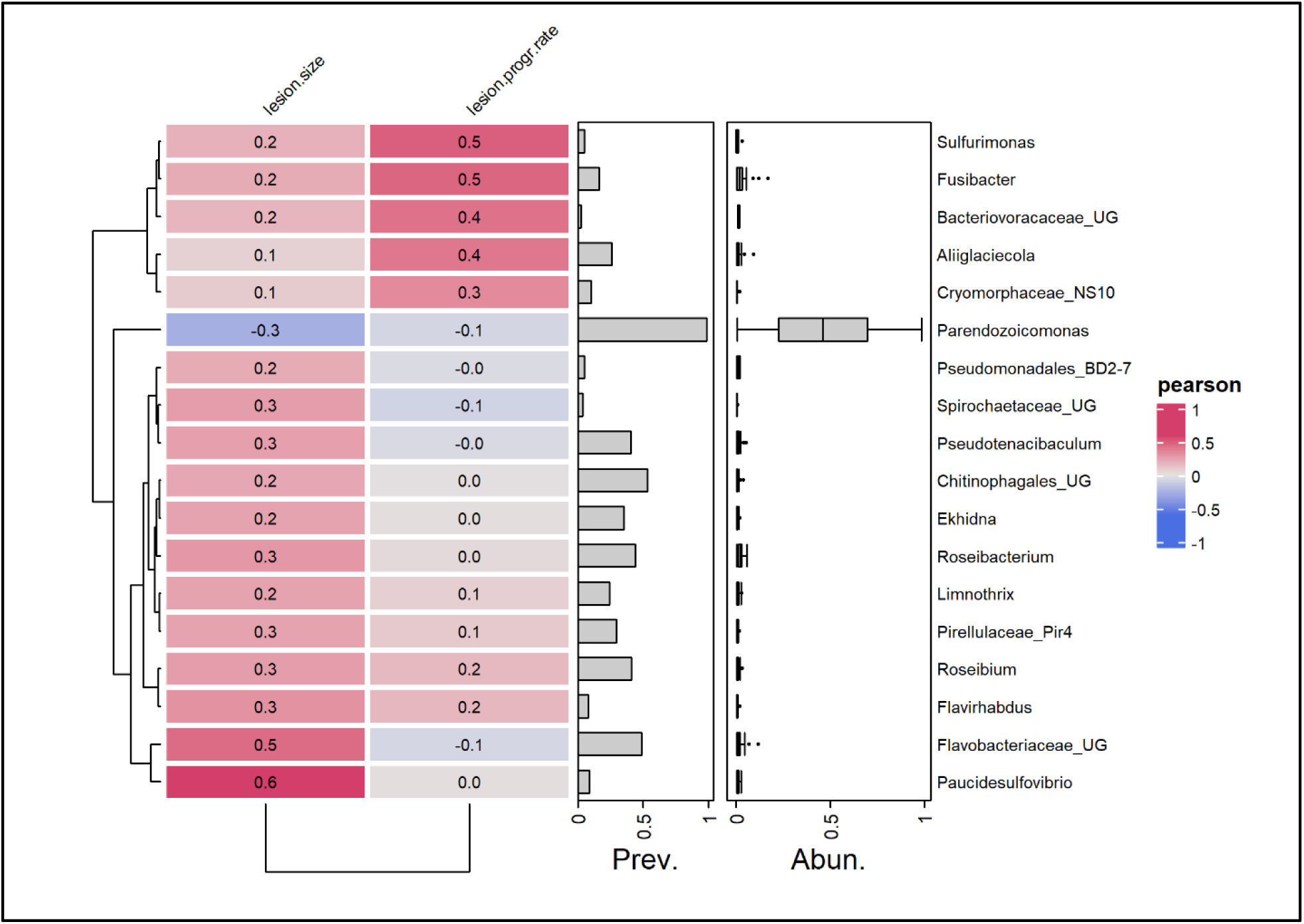
Eighteen genera from diseased tissue of *P. cylindrica* that showed Pearson’s correlation coefficient either above 0.3 or below −0.3 with lesion size or lesion progression rate. The bar graph shows genus prevalence (Prev.), and the box plot shows genus relative abundances (Abun.).

### 3. Environmental Monitoring

Chlorophyll a concentration ranged from 0.09 to 0.26 µg L^-1^ in Luminao seawater samples and from 0.13 to 2.09 µg L^-1^ in Tumon samples (Figure 12A). Chlorophyll a concentration differed significantly between the sites (ART, F = 46.8, p < 0.01) and time points (ART, F = 11.1, p < 0.01). Nitrate concentration ranged from 0.06 to 0.10 mg L^-1^ in Luminao samples and from 0.06 to 0.20 mg L^-1^ in Tumon Bay (Figure 12B), and these values were also significantly different between sites (ART, F = 53.7, p < 0.01) and among time points (ART, F = 53.7, p < 0.01). There was an interaction between site and time point for both the chlorophyll a (ART, F = 15.3, p < 0.01) and nitrate concentration (ART, F = 5.92, p < 0.01). The nitrate concentration in Tumon Bay peaked in May 2024 (Figure 2), coinciding with an extensive summer macroalgal bloom that covered many coral colonies, including those tagged for disease monitoring and microbial community sampling (Figure 12C-D).

**Figure 12.**
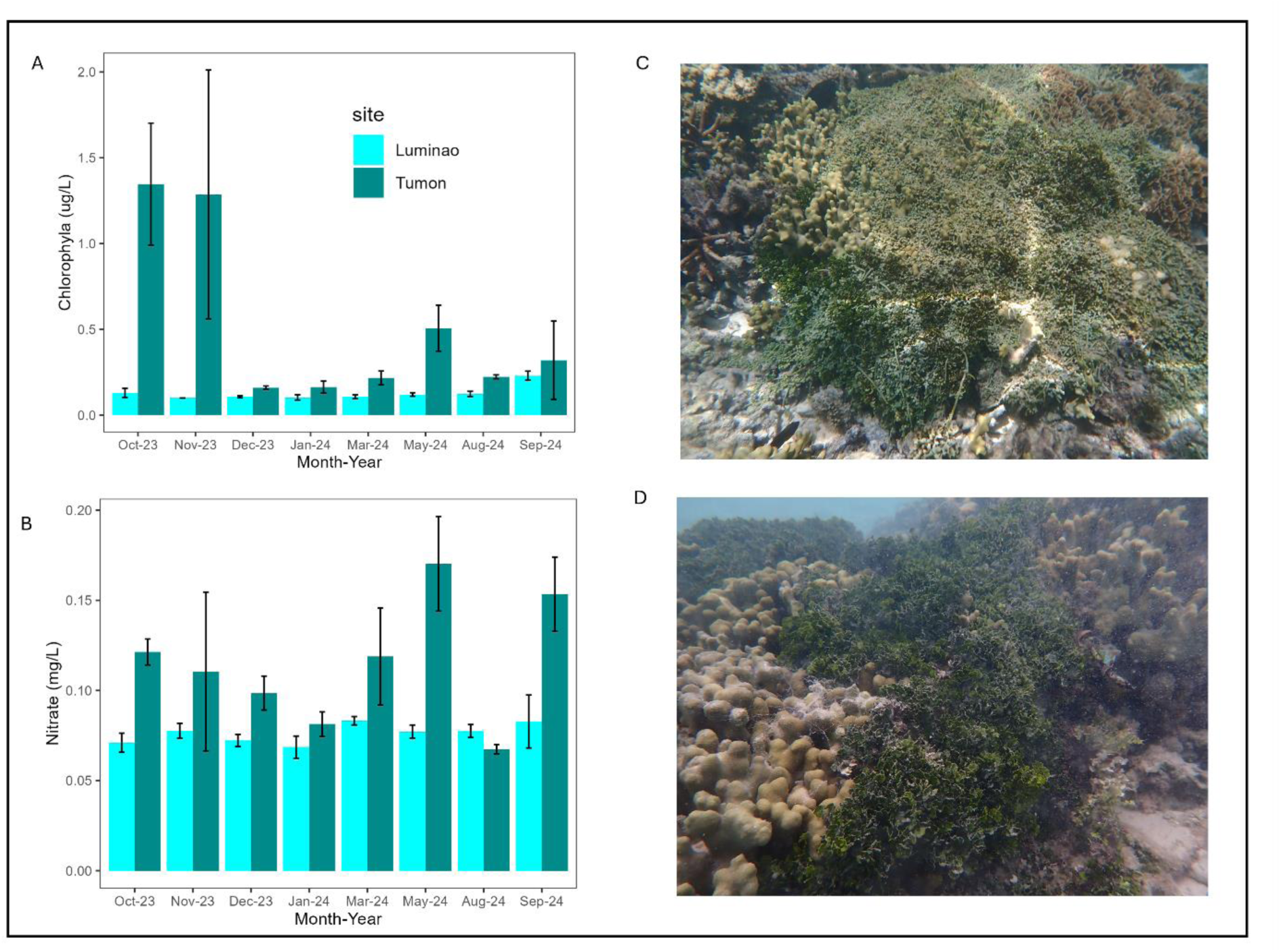
Chlorophyll-a (A) and nitrate (B) concentrations for two reef sites (Luminao and Tumon Bay) measured for a period of one year between October 2023 and September 2024. Photographs (C, D) of macroalgal bloom in Tumon Bay during summer 2024

Daily rainfall during the study period exhibited substantial variability, with a maximum of 141.5 mm on April 4, 2024. Monthly totals ranged from 41.2 mm in March 2024 to 494.5 mm in August 2023. During the rainy season, the monthly totals were always above 200 mm; the transition month between rainy and dry season could be distinguished by the monthly totals between 100-200 mm, and in the dry season, monthly totals were usually below 100 mm (except for an annual one-day rain in April 2024). The strong seasonal differences in rainfall were reflected across the sampling interval (Figure 13A).

**Figure 13.**
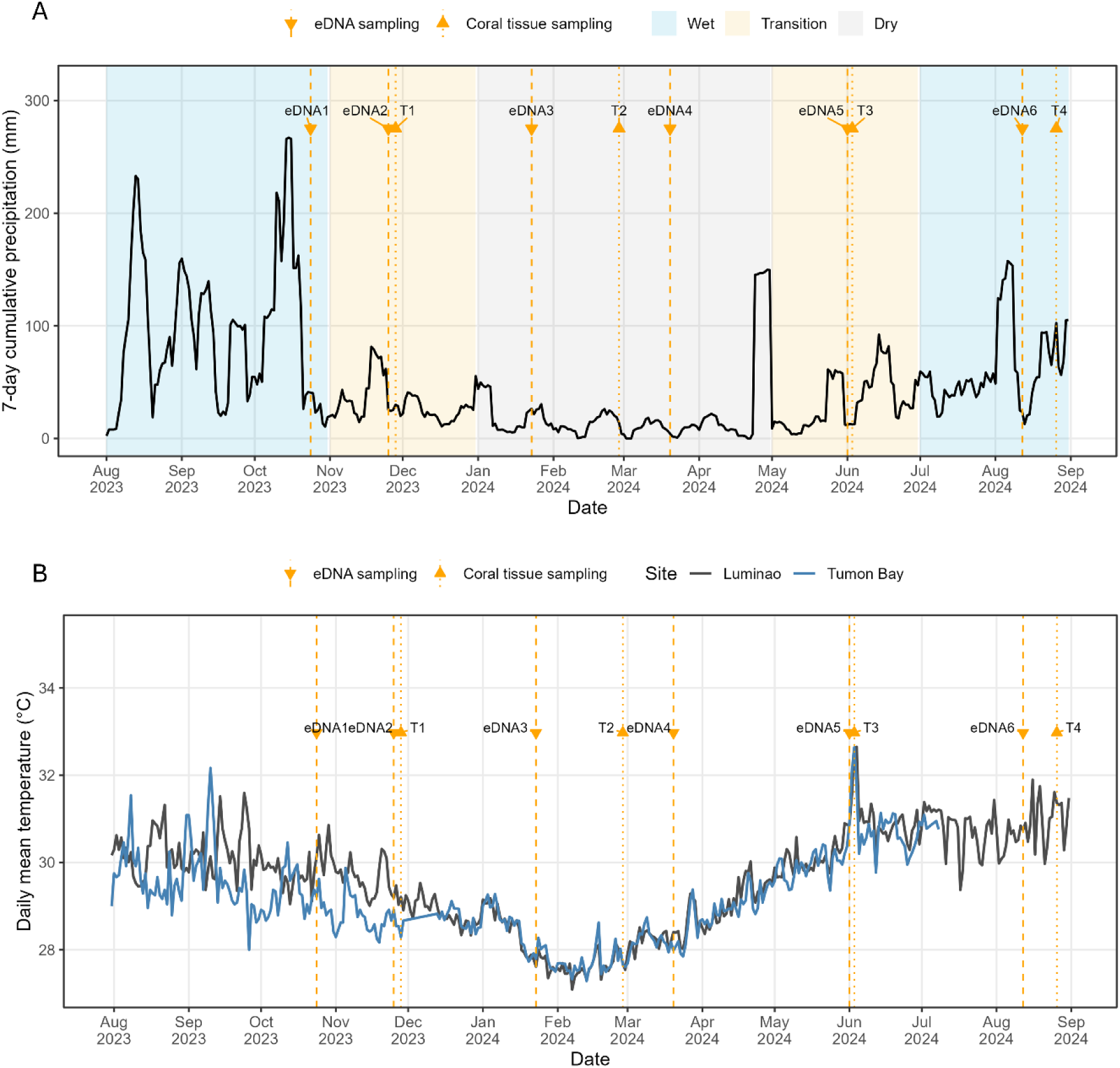
Temporal patterns in precipitation and seawater temperature at two reef sites on Guam between August 2023 and August 2024. **(A)** Seven-day cumulative precipitation (mm) from the Guam International Airport weather station, with shaded backgrounds indicating wet, transition, and dry seasons.**(B)** Daily mean seawater temperature (°C) at Luminao (black) and Tumon Bay (blue).Vertical dashed orange lines indicate sampling dates for seawater eDNA (eDNA1–eDNA6), and dotted orange lines indicate coral tissue sampling of *Porites cylindrica* (T1–T4).

Seawater temperatures at Luminao and Tumon showed a clear seasonal pattern from August 2023 through August 2024, with daily mean values ranging from 27.1°C to 32.6°C (Figure 13B). Daily mean temperatures were the highest in June 2024, reaching 32.65 ± 1.38°C at Luminao and 32.64 ± 1.14°C at Tumon, and were lowest in February 2024, 27.09 ± 0.93°C and 27.28 ± 0.54°C, respectively (Figure 13B, mean ± sd). Differences between sites remained small throughout the record (typically <0.2°C), indicating similar thermal regimes at both sites (Figure 13B).

## Discussion

In this study, we combined long-term disease monitoring, short-term lesion dynamics, microbial community profiling, and environmental measurements to investigate the drivers of *Porites cylindrica* white syndrome (PCYLWS) across two contrasting reef sites in Guam. PCYLWS prevalence and lesion dynamics differed markedly between sites, with higher long-term prevalence and faster lesion progression at Luminao despite smaller lesion sizes. Microbial communities were strongly structured by sample type, with pronounced shifts in community composition between healthy and diseased tissue and high temporal variability in lesion-associated microbiomes. Environmental microbial communities and nutrient conditions also varied across sites and seasons, suggesting potential links between environmental parameters and disease presentation. Together, these results indicate that PCYLWS is a dynamic condition shaped by both host-associated microbial changes and local environmental conditions.

To place these findings in an environmental context, this is the first study combining monitoring of eutrophication, environmental bacterial communities, and coral disease dynamics in Guam. Thus, it presents a unique perspective on seasonal and site-specific variations of ecological drivers and their potential links to the prevalence of white syndrome (WS) in the dominant reef-building coral *P. cylindrica.* Tumon Bay, which is chronically impacted by eutrophication due to high coastal development, heavy recreational use, groundwater seeps, and outdated sewage infrastructure (12), was confirmed to sustain elevated nitrogen levels year-round compared to the oligotrophic reference site of Luminao. Specifically, data from this study showed that 50% of nitrate concentration measurements (i.e., 4 out of 8 monthly surveys) in Tumon Bay exceeded the 0.1 mg L⁻¹ threshold for dissolved inorganic nitrogen (DIN), which was recently determined to be critical for coral reef health and biodiversity (42). Seasonal nitrate peaks were recorded in May and September 2024, coinciding with increased water temperatures and the annual bleaching season, resulting in an extensive macroalgal bloom that covered many coral colonies for several months. In parallel, chlorophyll concentration mirrored this enrichment pattern, with increased phytoplankton abundance at the beginning of the survey period (October–November 2023) and a smaller one in May 2024. These pulses likely reflect additional organic carbon input, potentially stimulating microbial activity (16). In contrast, Luminao showed consistently low nitrate and chlorophyll levels throughout the year, confirming its oligotrophic status.

Nutrient enrichment is well known to promote phytoplankton and macroalgal blooms and to increase dissolved organic carbon availability, typically associated with increases in bacterial biomass and shifts in bacterial community assemblages, i.e., microbialization (8, 9). As such, eutrophic conditions are often associated with increased prevalence and severity of coral disease (10–13). Based on these patterns, we expected Tumon Bay to present a higher prevalence or severity of white syndrome (WS) tissue loss disease in *P. cylindrica*, a species known to be both abundant and chronically susceptible to WS in Guam (20, 43). Surprisingly, long-term monitoring data since 2011 revealed a persistent and higher prevalence of WS in Luminao at the oligotrophic site. Specifically, many large colonies, exceeding 1-2 meters in diameter and exhibiting high WS prevalence, have been observed in Luminao but not in Tumon Bay, thereby contributing to the overall differences in WS prevalence between the sites. On the one hand, WS lesions were larger and more rapidly colonized by turf algae at Tumon Bay, whereas the small lesions in Luminao showed a higher progression rate, but only for a short period, before reaching smaller sizes. These patterns highlight that coral disease dynamics are not solely driven by nutrient levels but also by site-specific differences in host demographics (i.e., abundance, size, and age), host susceptibility, pathogen reservoirs, and microbial community composition, both in the surrounding environment and within the coral holobionts.

Both study sites exhibited significant variation in their water-column bacterial community structures, with higher relative abundances of Flavobacteriales, Rhodobacterales, and Enterobacterales in the nitrogen-enriched waters of Tumon Bay. This pattern is consistent with previous studies showing that copiotrophic bacterial groups are more abundant in organic matter-rich water and sediment (16, 44), as they play key roles in decomposing large organic molecules (e.g., complex polysaccharides) and thrive on marine detritus (45, 46). In addition, Enterobacterales showed a higher relative abundance in the Tumon sediment sample than in Luminao. Many members of Enterobacteriales are known animal and plant pathogens, including coral pathogens (47). This order is also known to increase along eutrophication gradients (15, 44, 48). It should be noted, however, that the elevated relative abundance of Enterobacteriales in seawater and sediment in Tumon was only observed from October to November 2023; the remaining time points showed values similar to those in Luminao. That increase was most likely related to the Oct-Nov 2023 sampling period, which followed a few months of the rainy season, during which additional nutrients and sediment are transported to coastal waters.

Establishing the causation of coral diseases presents numerous challenges and is rarely achieved (49–51). One key difficulty lies in the complexity of the coral holobiont, which inhabits seawater containing approximately one million bacteria per liter (52). Any tissue injury, such as a fish bite or physical abrasion, can attract opportunistic bacteria to the lesion, thereby further accelerating tissue degradation (53, 54). Thus, the bacterial community in coral lesions is highly complex and may obscure the presence of the primary pathogen. In the current study, multiple bacterial groups (six classes, 13 orders, and 17 families) were significantly more abundant in diseased tissue than in healthy tissue. This highlights the bacterial complexity of lesions, likely restructured by the settlement of opportunistic decomposers of necrotic coral tissue. Moreover, the composition of the lesion-associated bacterial community varied significantly across seasons and locations, indicating that environmental parameters influence the process of opportunistic colonization. Due to the scarcity of consistent associations between specific pathogens and particular coral diseases, scientists propose that the “one pathogen: one disease” paradigm may not apply to coral diseases, and alternative etiologies must be considered (47, 55).

One recognized hypothesis is that disease arises from a disruption in the coral’s ability to maintain a stable beneficial microbiome in response to environmental stress, chemical toxicity, or viral infection (47). This may compromise the host’s capacity to curate its microbiome, leading to the loss of coral-beneficial symbionts and the emergence of microbial dysbiosis. The state of microbial dysbiosis has been associated with multiple coral diseases (56–58). In this study, the microbiome community compositions of PCYLWS lesions versus healthy tissues exhibited key characteristics of microbial dysbiosis. The healthy microbiome was dominated almost entirely by the genus *Parendozoicomonas*, whereas the microbiome in PCYLWS lesions exhibited significant spatial and temporal variability. No single bacterial genus was detected across all 156 diseased samples. Instead, various bacterial genera were observed at varying frequencies, with the highest prevalence recorded for *Ruegeria* (90%), followed by *Muricauda* (85%). Members of the family *Paracoccaceae* (formerly *Rhodobacteraceae*), including *Ruegeria* and many other undescribed taxa, are commonly found in healthy coral mucus and skeleton (59, 60), but they flourish in diseased tissues (61, 62). Interestingly, some mucus-associated bacteria, such as *Ruegeria* strains, exhibit antimicrobial activity against other bacteria, including *V. coralliilyticus*, and have been explored as potential coral probiotics (63, 64). These findings suggest that *Paracoccaceae* may play dual roles in coral health and disease, depending on environmental context and host conditions.

In contrast, the genus *Parendozoicomonas*, which was dominant in healthy tissue, is closely related to *Endozoicomonas* and was described more recently from sponges (65, 66). *Parendozoicomans* is the primary bacterial genus found in healthy *Porites* spp. in Guam (67) The genus name is based on the suggested *Candidatus Parendozoicomonas poriteae,* proposed based on the metagenomically assembled genomes (MAGs) from healthy *Porites* spp. found across the Pacific (68). Members of the family *Endozoicomonadaceae* are known to be the most abundant bacterial group in many healthy corals, and growing evidence suggests they contribute to corals’ health and function. For example, their genomes encode biosynthesis pathways for vitamins (B1, B2, B7, and B9), which are essential to corals and *Symbiodiniaceae*; however, neither corals nor *Symbiodiniaceae* can synthesize these vitamins (69, 70). In addition, members of *Endozoicomonadaceae*, including *Ca. P. poriteae* can metabolize dimethyl sulfoniopropionate (DMSP), thus helping the coral holobiont with organic matter recycling (68, 71).

Although the microbial dysbiosis model accounts for shifts in coral-associated microbiomes, it does not explain the frequent occurrence of transmissible lesions in coral diseases. It has previously been demonstrated that WS lesions in *P. cylindrica* can be transmitted between individuals, suggesting that at least some lesions may involve infectious agents (72). If viral infections trigger dysbiosis, transmission may still be possible. However, the limited current research on coral viruses prevents us from concluding or even speculating about that possibility. The current funding is also consistent with the presence of multiple etiologies of white syndromes (21). It remains possible that coral diseases with similar symptoms (e.g., white syndromes) in the same host species can be caused by a variety of bacterial species, whether originating from sediment, seawater, or coral commensals. There is already some evidence for such multi-causations: WS in *Pocillopora damicornis* has been linked to infections by *Vibrio coralliilyticus* during coral immunosuppression caused by increased seawater temperature (73), as well as by *V. harveyi* (74). Similarly, a combination of six different *Vibrio* strains can cause WS in three distinct coral species from distant Pacific locations (75), and both *V. coralliilyticus* and *V. owensii* induce tissue loss in *Montipora capitata* (76, 77). As those examples demonstrate, multiple causations exist among different species within a single genus; therefore, the hypothesis that bacteria of different genera can cause WS in the same coral host cannot be excluded. Specifically, since bacteria other than *Vibrio* are rarely investigated for their role in coral pathogenicity, they are more difficult to isolate than *Vibrio* spp. PCYLWS lesions examined in this study may have been initiated by a variety of bacteria, such as *Maricauda*, *Ruegeria*, *Aligola*, or *Vibrio*, acting alone or in combination. It should also be noted that *P. cylindrica* from both sites harbors highly stable Symbiodiniaceae algal communities across time and space, specifically *Cladocopium* C15-ITS2 type lineages. This suggests that shifts in algal symbionts are unlikely to account for the observed differences in disease presentation or causation from the two studied sites (78).

Correlation analyses in this study suggest that the genera *Sulfurimonas* and *Fusibacter* might contribute significantly to lesion progression, as both showed positive associations with lesion progression rates. *Fusibacter* has also been detected multiple times in the pathobiomes of Stony Coral Tissue Loss Disease (79–81). In addition, a correlation between Flavobacteriaceae relative abundance and lesion size suggests that this family may exhibit opportunistic colonization and tissue degradation. These associations underscore the importance of further investigating these bacteria. Significantly, compositional methods derived from V4-16S rRNA sequencing may underestimate the accuracy of correlations between abundance and lesion severity (e.g., the presence and abundance of one bacterium can substantially influence the detection and abundance of other bacteria). Complementary methods targeting absolute abundance (e.g., qPCR) would likely provide more precise insights into bacterial contributions to tissue loss.

In conclusion, this is the first study to investigate the bacterial dynamics of white syndromes in *P. cylindrica* in Guam and the influence of eutrophication on the associated microbiomes. Lesion sizes were larger at the eutrophic location, while disease prevalence and activity were highest where host density was greatest, suggesting that multiple factors drive disease dynamics. Copiotrophic bacteria (Flavobacteriales, Rhodobacterales, and Enterobacterales) were enriched in high-nitrate waters, potentially contributing to lesion size in coral found at Tumon Bay. In contrast, *Vibrio* species were more abundant at the low-nitrate site with higher WS prevalence and activity. Diseased coral tissues harbored a taxonomically diverse bacterial consortia, complicating the identification of a single causative agent (a bacterium or even a single genus). However, the frequent detection of *Ruegeria*, *Maricauda*, and *Vibrio* in lesions suggests their possible causation or contribution to the disease process; however, this could only be verified experimentally. These findings support the view that coral diseases may arise from context-dependent interactions among multiple opportunistic pathogens. Future work should also include additional functional microbial groups, such as viruses, and functional approaches to fully understand coral pathobiomes. This study provides a comprehensive baseline on the bacterial dynamics associated with PCYLWS. It lays the foundation for future research into the etiology of this disease, which affects a key reef-building species in the Guam region.

## Data Accessibility

The raw reads have been deposited into the NCBI SRA database under the BioProject accession: PRJNA1227205 and BioSample accession numbers: SAMN46988364 - SAMN46988559.

## Acknowledgments

We want to thank Jon Rubin and Zoe Ariellius for their assistance with sample collection in the field. Coral collection was possible thanks to the permit number: SCR-MPA-23-014 issued by the Division of Aquatic Wildlife Resources (DAWR) of the Department of Agriculture (DOAG).

